# ll-optical Diamond Heater-Thermometer enables versatile and reliable thermal modulation of ion channels at the single-cell level

**DOI:** 10.1101/2025.06.03.657561

**Authors:** J.-S. Rougier, E. Glushkov, S. Guichard, J. P. Kucera, V. Zeeb, H. Abriel

## Abstract

A living cell is a nonequilibrium thermodynamic system where, nevertheless, a notion of local equilibria exists. This notion applies to all micro- and nanoscale aqueous volumes, each containing a large number of molecules. This allows one to define sets of local conditions, including thermodynamic ones; for instance, a defined temperature requires thermodynamic equilibrium by definition. Once such a condition is fulfilled, one can control local variables and their gradients to theoretically describe the thermodynamic state of living systems at the micro- and nanoscale. Performing ultralocal experimental manipulations has become possible thanks to the patch clamp technique to control the cell membrane potential, as well as fluorescent imaging to monitor molecular concentrations and their intracellular gradients. However, precise temperature gradient control at the micro-/nanoscale has yet lacked a reliable experimental realization in a living cell.

Here, we present a new methodology – microscale control of a temperature gradient profile in aqueous media by a fully optical Diamond Heater-Thermometer in a plug-and-play configuration combined with the patch clamp technique. In particular, we demonstrate applications of the combined Diamond Heater-Thermometer-patch clamp approach for the fast and reproducible thermal modulation of ionic current from voltage-gated Na_v_1.5 sodium channels expressed in HEK293 cells and freshly isolated ventricular mouse cardiomyocytes. Such an approach of manipulating the ultra-local temperature down to the nanoscale has the potential to uncover previously inaccessible phenomena in various physiological intracellular processes related to the endogenous nanoscale heat sources, such as open ion channels capable of producing Joule heat.

## Introduction

Studies on the temperature effects on single cells, tissues, and the whole body show its multifarious role at all structural levels in living organisms [1], [2], [3], [4]. The most intriguing are the thermal effects related to intracellular calcium signaling [5], [6], [7], [8], [9], thermally induced cell morphology changes [10], [11], [12], manipulating embryogenesis via nanoscale temperature control [13], thermal activation of single kinesin molecules [14], and thermal activation of ThermoTRP channels [15]. The exceptional importance of temperature as a thermodynamic parameter in life is illustrated by the small range of temperature where life forms can exist – just around one hundred degrees Celsius, in comparison with the physical range from 0 Kelvin (K) to Plank’s temperature of 10^32^ K [16]. An even smaller temperature range of 37–42 °C was recently identified as the most energy-efficient for processing neural signals [17]. The existence of life in such a narrow temperature range puzzlingly coincides with experimental data demonstrating the possibility of artificially creating steep (up to 20°C/μm) steady-state temperature gradients in volumes of aqueous media at the micro-/nanoscale [18], [19]. This corresponds to the extremely low thermal conductivity of water [20] under the assumption of Fourier heat transfer phenomena by conduction [16], [21].

Experimental progress during the last five decades enabled ultra-local intracellular control of the membrane potential in single cells by the patch clamp technique [22], and the measurement of ionic/molecular concentrations and their gradients by fluorescent imaging using specific fluorescent dyes [23]. However, the temperature gradient control at the micro/nanoscale has lacked a reliable experimental implementation in a living cell, particularly regarding ion channels. Ion channels allow the movement of ions through an ion-impermeable lipid bilayer and are crucial for many biological events. As with other proteins, temperature dramatically influences the biophysical properties of all ion channels [24], [25], [26]. The patch clamp technique allowed the investigation of those biophysical properties (e.g., current amplitude, channel kinetics) [27]. Many of those investigations are performed at non-physiological temperatures (22-23°C) for technical purposes (e.g., ensuring the stability of the recording as the fluidity of the lipid bilayer increases with temperature). Although heating devices (e.g., chamber perfusion using the Peltier approach [28], [29], [30]) have been developed to investigate ion channels at physiological temperatures, their main limitations are 1) the absence of local and precise increase of temperature leaving the membrane integrity of cells surrounding the investigated one ‘intact’ for further recordings; 2) the inability of increasing the temperature within milliseconds does not enable the investigation of biophysical properties of ‘fast ion channels’, such as the voltage-gated sodium channels family; 3) the unreliability of the recorded parameters due to the methods’ inability to apply different temperatures at different times to the same cell (due to the relaxation (millisecond time scale) to achieve a thermodynamic equilibrium).

Here, we present a new reliable methodology to control the temperature gradient profile in aqueous media at the microscale with an all-optical Diamond HeaterThermometer (DHT), described previously in [19], in combination with patch clamp whole-cell electrical recordings. Briefly, a small fluorescent diamond particle is embedded into the tip of a pulled glass microcapillary pipette. The diamond particle is connected to a tapered optical fiber guiding both the excitation light and fluorescence (produced by the temperature-sensitive color centers inside the diamond). The particle has a graphite shell that efficiently absorbs laser light, enabling it to function as a calibrated thermometer and an ultra-local precision heat source [19]. The DHT approach of manipulating as well as sensing the local intracellular temperatures down to the nanoscale, combined with the patch clamp technique, has the potential to control previously inaccessible intrinsic nano-/microscale energy transformation effects in various physiological intracellular processes (including structural, electrical, signaling, and biochemical processes) related to the endogenous nanoscale heat sources, such as the predicted heat release from the open ion channels [31], ionic pumps [32], mitochondria [33], calcium release processes [6], [7], and muscle contraction [34], [35].

The simplicity of the DHT in the fiber-coupled configuration ensures its compatibility with most existing laboratory setups, without requiring any modifications to the optical paths. The solid-state nature of the DHT guarantees the absence of photobleaching, allowing long experimental protocols of thermal stimulation applied to individual living cells. Being fully optical, the method does not produce electromagnetic noise hindering patch clamp recordings. Moreover, due to the ultra-local heating with millisecond time-scale stabilization of the temperature gradient/clamp profile and precise nanoscale 3D positioning of the diamond, this method overcomes the thermal expansion issue associated with conventional slow heating of the experimental chamber, which shifts the specimen out of the focal plane. Other advantages of the DHT technique include its infinite photostability and environmental insensitivity, providing greater flexibility of thermal stimulation and control compared to existing techniques. This set of capabilities can open new directions in cellular physiology, such as deciphering ultra-local thermodynamic phenomena of intracellular processes and their influence on cellular function.

## Experimental results

The experiments described here were performed using the experimental setup shown in Fig. 1a. It consisted of an inverted optical microscope, a patch clamp module, and a DHT device (more in Materials & Methods). HEK293 cells stably expressing the human voltage-gated sodium channel Na_v_1.5 were patch clamped (Fig. 1b), and the sodium current was recorded. During the recording, heat pulses of 30 seconds were applied to the tip of the fiber-coupled DHT pipette to modulate the biophysical properties of the sodium current (peak current and decay time) as depicted in Fig. 1d. Similar experiments were performed in HEK293 cells not expressing Na_v_1.5 channels (wild-type HEK293 cells) as a negative control (Fig. 1c).

**Figure 1.**
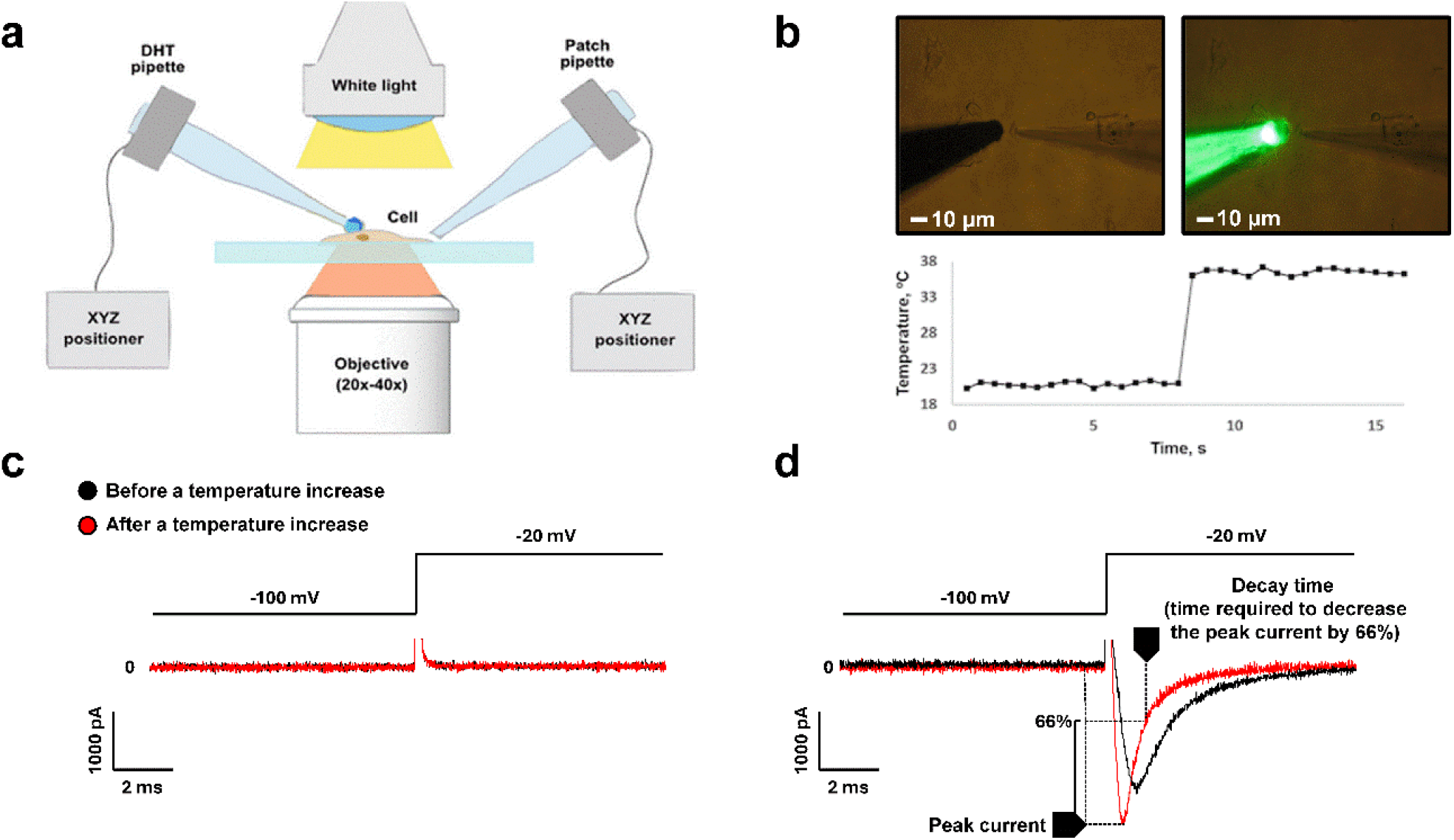
a) Schematic representation of a combined DHT-patch clamp setup. b) Microscope image of a clamped HEK293 cell with an approaching DHT pipette. c) Whole-cell patch clamp recording of the ionic current through the cell membrane in the wild-type cells and d) in cells stably expressing the voltage-gated sodium channel Na_v_1.5. The artefact of stimulation observed when modifying the membrane voltage has been cut for space purposes.

We then varied the laser power delivered to the DHT pipette to obtain heat pulses reaching various peak temperatures of DHT and consequently producing different steady-state temperature profiles in the cell (Fig. 2). The slope of the temperature profile depends on the laser power delivered to the heat source, its size, and the distance between DHT and the cell (in general around 5-10 micrometers), as discussed in [18], [21]. With the increased temperature, as anticipated, the sodium peak current values increased, and the current decay times decreased with the increased temperature (Fig. 2a and b). Such effects were observed only in cells stably expressing the voltage-gated sodium channel Na_v_1.5 and not in the wild-type (see Fig. S1). It is important to note that the laser illumination had little to no effect on those biophysical properties (Fig. S2). Only the laser light absorbed by the diamond particle’s surface at the tip of the DHT pipette, producing local heat, played an observable role. The exact temperature of the heat pulse can be measured in real-time, which is one of the key advantages of the DHT system compared to a more standard direct laser heating. Another advantage is the reduced photodamage to living cells, as only several milliwatts of laser power are sufficient to execute thermal stimulation compared to hundreds of milliwatts required by a conventional infrared (IR) laser heating approach.

**Figure 2.**
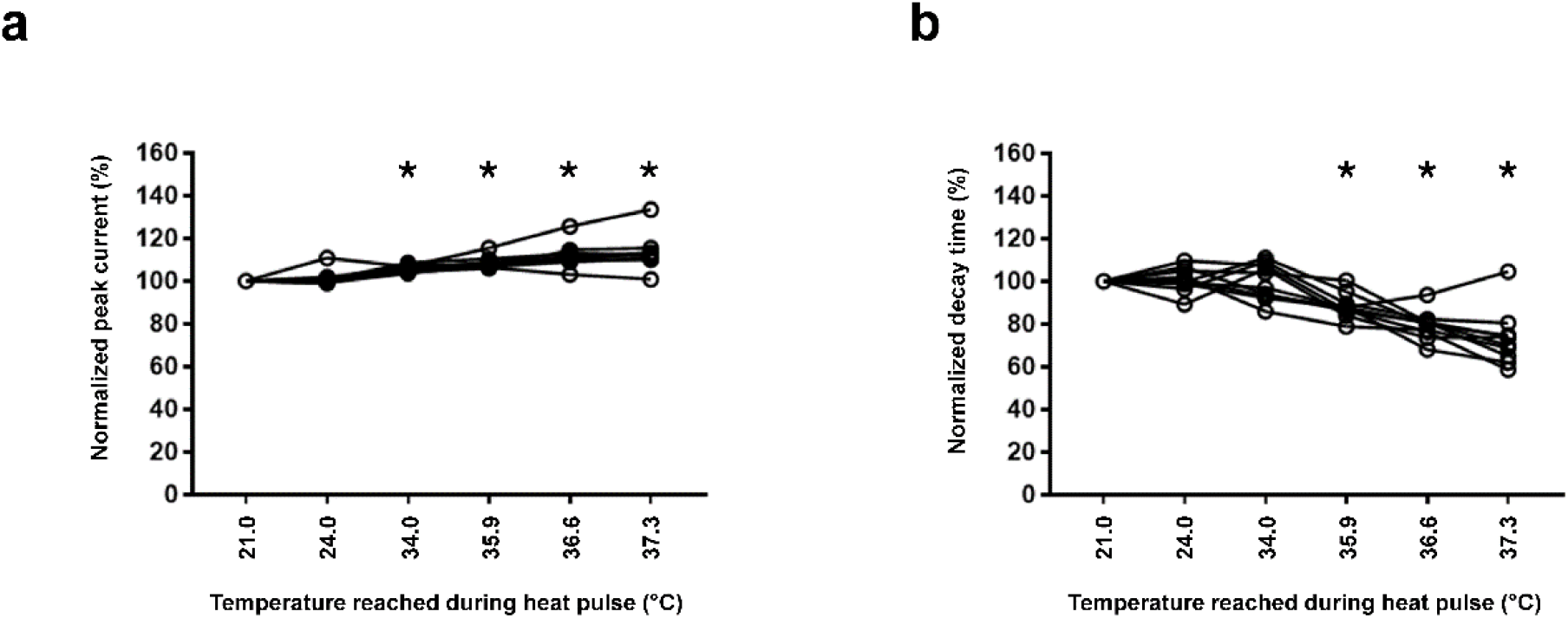
Effect of local heat pulse from the DHT. a) Peak current increased with the increased temperature of the pulse. b) Decay time decreased when the temperature of the pulse increased. These effects are observed only in cells stably expressing the voltage-gated sodium channel Na_v_1.5 and not in wild-type cells. *n* =10 cells. *; *p* < 0.05 versus 21.0°C.Each line, connecting a circle, corresponds to the modification of the parameter for different temperatures applied to a single cell.

To further test the reliability and repeatability of the DHT device in modulating the ionic current, we performed active thermal cycling between various temperatures (Fig. 3). Within 15 seconds after modulating the temperature from 19 °C to 36 °C, we observed that the normalized average peak current increased by ~15% (Figure 3a). An inverse effect of ~20% change is observed with a similar heat pulse on the average normalized decay time (Figure 3b). Interestingly, this modulation is reproducible, highlighting the absence of harmful effects of this approach on the cell (Fig. 3c and 3d).

**Figure 3.**
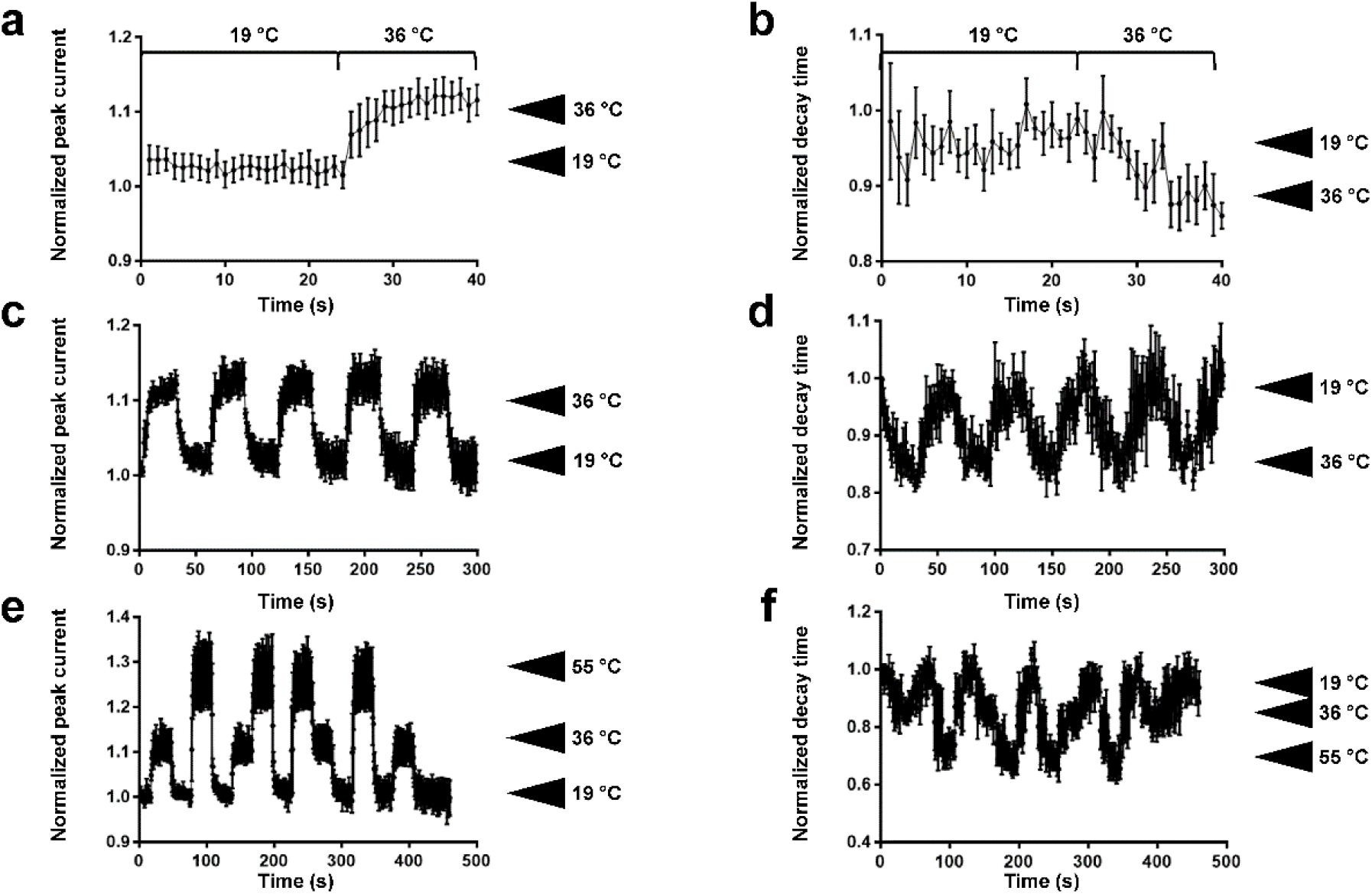
Repeatability and reproducibility of the thermal modulation of sodium current. a) Normalized average peak current (*n* = 6) and b) normalized average decay time (*n* = 5) modulation by a heat pulse from 19 °C to 36 °C. c and d) Repeatability of the thermal modulation of the peak current (c) and decay time (d). Normalized average peak current (e) and decay time (f) modulation (*n* = 3) by different temperatures (36 °C and 55 °C), showing the reproducibility of this modulation independently of the sequence of temperature steps applied. The increased noise observed at the end of the decay time measurements (d) is related to the instability of the seal.

One of the key features of the DHT device is its ability to selectively apply an ultra-local temperature profile owing to its microscopic size. We therefore decided to apply different temperatures (36 °C and 55 °C) in various orders to modulate the peak current. This is hardly feasible with a classical approach due to the time required for the thermal dissipation after switching to a new temperature. As shown in Figure 3, the peak current increase and the decay time decrease observed at 36 °C and 55 °C are accurately and reproducibly observed independently of the order in which those temperatures were applied. This indicates that the DHT device induced a rapid temperature change due to its ultralocal application and a rapid thermal relaxation (millisecond time scale) (Fig. 3e and 3f). Thanks to these properties, the heat pulse applied to the cell can be relatively short, and increasing the temperature to 55°C for a short time to record the sodium current doesn’t affect the cell viability.

In parallel to experiments with HEK cells, we obtained freshly isolated adult ventricular mouse cardiomyocytes, which are more temperature-sensitive than HEK cells, and recorded sodium current from their endogenously expressed Na_v_1.5 channels. We observed similar thermal modulations on the peak sodium current and the decay time in cardiac cells without damaging the surrounding ones (Fig. 4a and 4b). The increase in peak current amplitude in cardiomyocytes is significantly higher than in HEK293 cells expressing Na_v_1.5 channels due to the higher sensitivity to temperature of these cardiac cells.

**Figure 4.**
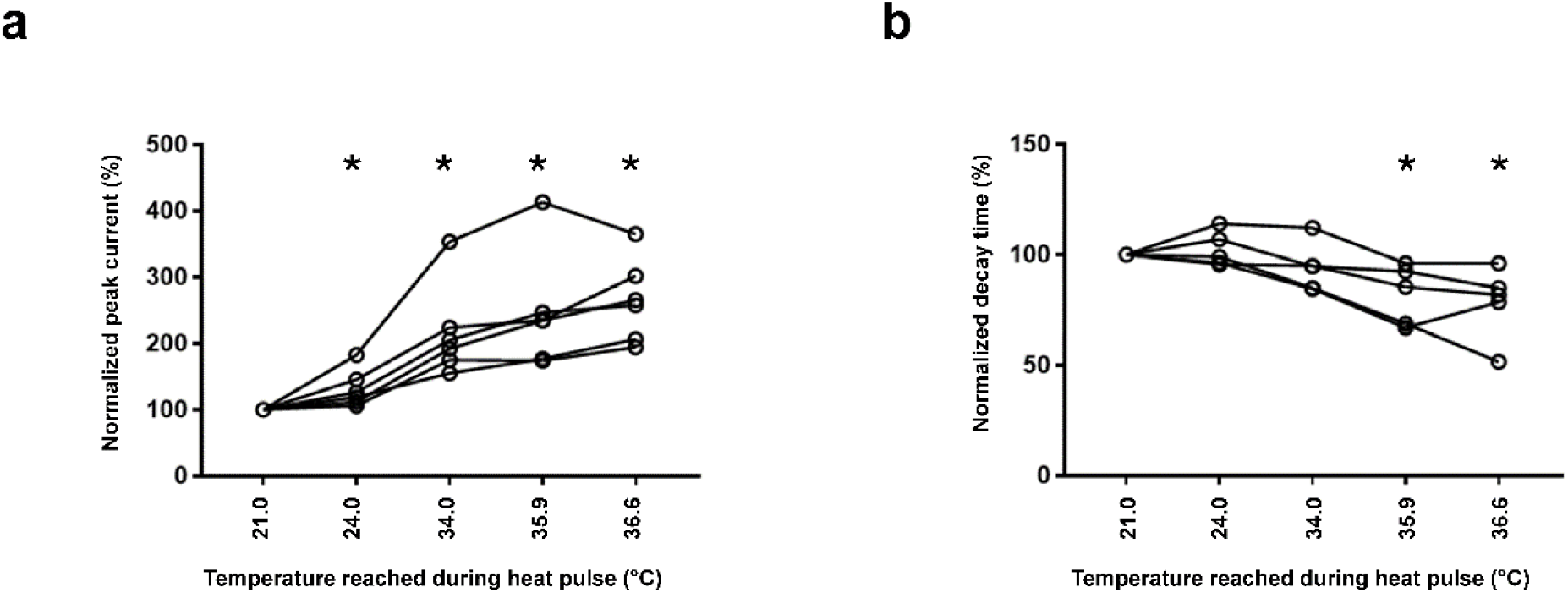
Effect of local heat pulse from the DHT on the peak sodium current (a) and decay time (b) extracted from the corresponding patch clamp recordings using freshly isolated murine adult ventricular cardiomyocytes. *N* = 5-6 cells. *; *p* < 0.05 versus 21.0 °C. Each line, connecting a circle, corresponds to the modification of the parameter for different temperatures applied to a single cell.

Then, cardiac action potentials were recorded before and during a heat pulse to assess the effect of heat on different parameters (Fig.5a and 5b). As observed in Figure 5a and represented in Figures 5c and 5d, a heat pulse induced a significant increase in the time to maximum amplitude and the APD90 (Fig. 5). In addition, the upstroke velocity was increased, which reflects the increase of sodium current observed in Figure 4a (upstroke velocity in mV/ms at 21.0 °C: 81 ± 4; at 35.9 °C: 97 ± 4; *n* = 8; *p* < 0.05).

**Figure 5.**
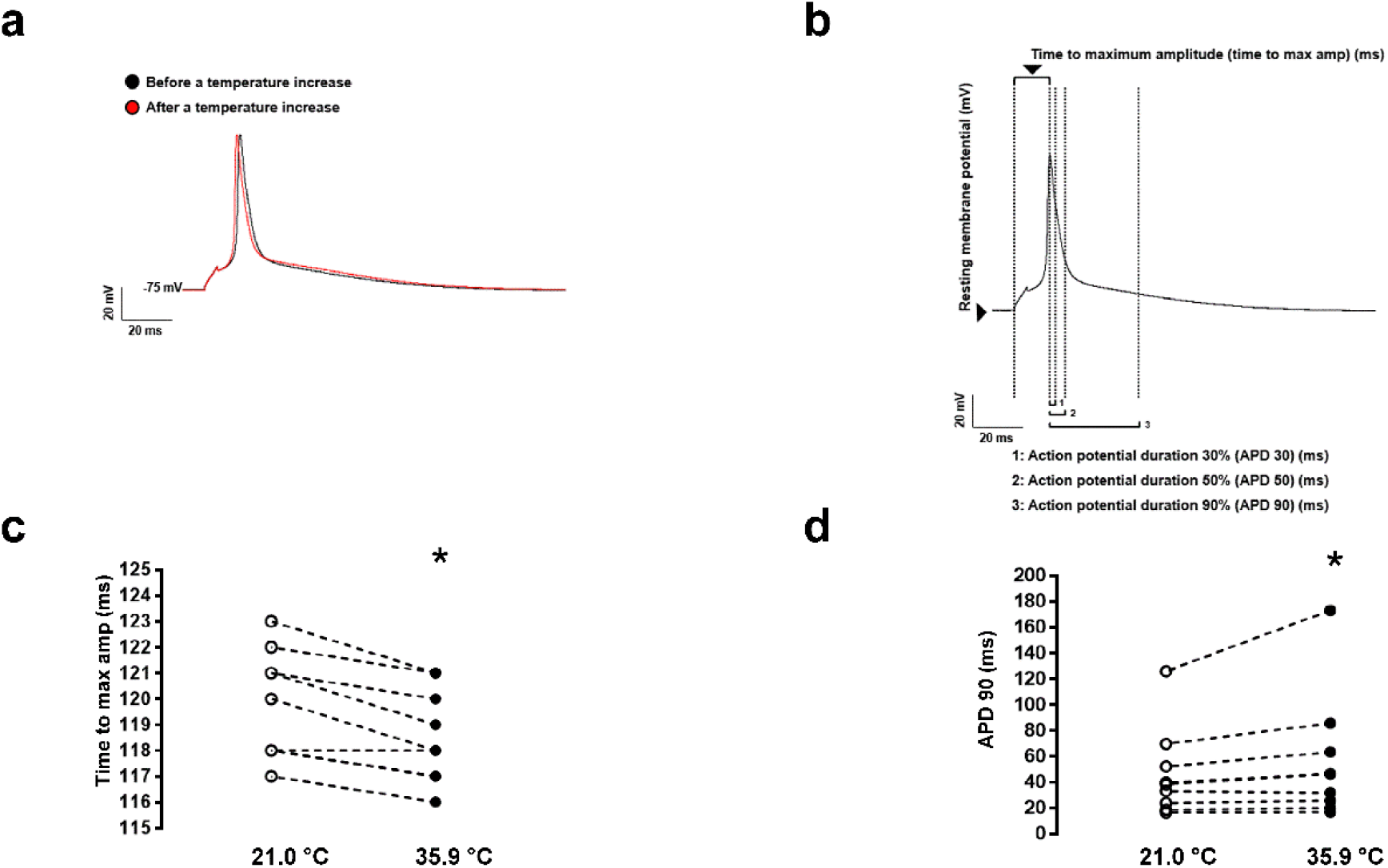
Murine adult ventricular cardiac action potential recordings showing the effect of a heat pulse (a) and the different biophysical parameters investigated (b). Only the time to maximum amplitude (max amp; c) and the 90% action potential duration (APD90; d) are significantly different comparing the room temperature (white circle, t° = 21.0 °C) to heat pulse conditions (black circles, t° = 35.9 °C; *n* = 6 cells; *; *p* < 0.05 versus 21.0 °C).

## Discussion

### Comparison of published ion channel data and implications for future experiments

Our group investigated the modulation of the biophysical properties of Na_v_1.5 wild-type channels by increasing the temperature several years ago using a similar approach (patch-clamp setup and cell line), but with a heating perfusion system instead of the DHT technique [30]. No significant difference was observed between our previous data and the data presented in this manuscript regarding the effect on the peak current measured using the whole cell configuration. However, the decrease in decay time seems smaller in this study than in the one using the heating perfusion system, which suggests that temperature was partially heterogeneous [30].

The time required in our previous publication to reach thermal stability and perform the different recordings was approximately 1 minute [30], whereas with the DHT approach, this value is closer to 15-25 seconds. One explanation for this difference may be the position of the heating source. Heating perfusion systems are generally challenging to position close to the cell, as the perfusion of the solution damages the cells (stretching forces). In contrast, the DHT may be more closely positioned to the cell without affecting its viability and, therefore, reduce the time required to heat the cell to the desired temperature. In addition, the time required to reach the requested temperature at the tip of the DHT probe is in the range of milliseconds [36], which is significantly shorter than the time to measure to obtain the stability of the current (~seconds). This delay can be explained by the time required to warm the entire water environment surrounding the cell, and is also observed when using a Peltier device to heat a water bath. Such a delay may be problematic for investigating the limits of temperature-dependent biophysical modulations that occur at a sub-second timescale. The ease of positioning the DHT close to the target, without affecting the stability of the seal or the noise, as in the case with the heating perfusion system, may be useful for investigating those parameters not only in a whole-cell configuration but also in a single-channel recording experiment using the inside-out and outside-out configurations. Such experiments can provide more insight into the bandwidth of thermal modulation in ion channels.

Recording biophysical properties of ion channels at physiological temperatures or higher is a ‘‘must-have”. A plethora of investigations have highlighted the consequences of hyperthermia in organ dysfunction [6], [30] such as in the heart (the Brugada syndrome, heat stroke, and heat exhaustion), and the brain (neurological and cognitive alteration), which are probably due to the alteration of ion channel function at high temperatures. For example, ion channels from the TRP family show an increase in the current density by a factor of 8 with a temperature increase of 10°C [37]. Nevertheless, such recordings face multiple technical challenges. The ultra-local temperature controller presented in this manuscript will address many of these challenges and open up new avenues for investigating local temperature fluctuations and their implications for ion channel function.

### Ultra-local microscale thermodynamics

The precise control of temperature as an ultra-local variable assigned to the different micro-/nanoscale volumes in aqueous media in such a nonequilibrium thermodynamic system as the living cell should rely on the notion of local equilibria [16], [21], [38]. Fulfilling this notion ebables researchers to apply the temperature gradient correctly (thermodynamic force) at the micro-/nanoscale when studying the physiology of living cells. More than ninety years have passed since Lars Onsager discovered reciprocal relations between different forces and flows as a consequence of the time reversibility of microscopic dynamics in nonequilibrium systems where the notion of local equilibria exists (Nobel Prize 1968; [39], [40]). These relations are now close to being named the Fourth Law of Thermodynamics due to their universal fundamental applicability in nonequilibrium thermodynamic systems, including living cells. The existence of local equilibria in such systems allows one to introduce sets of local variables, including a thermodynamic one, such as temperature. The use of these intracellular variables and their gradients (forces), such as temperature gradient, voltage gradient, chemical potential gradient, and their corresponding flows – heat flow (Fourier’ heat conduction law), electrical current (Ohm’s law), diffusional flow (Fick’s law), to describe theoretically and implement experimental manipulations at the nano/micro scale. Each force is not an independent player, giving birth to a corresponding flow. Onsager’s law mutually intertwines all the forces through the interplay between flows induced by corresponding force-related processes, such as the conduction of heat, electrical currents, and diffusion. Classical examples of pair-reciprocal phenomena in physics are Peltier and Seebeck, or Dufour and Soret [40]. In general, Onsager’s theory can treat more than two thermodynamic forces at once, meaning that the introduction of nano-/microscale temperature clamp/control method in aqueous media enabled by the DHT can uncover the previously hidden microscale effects of energy transformation processes in living cells bound to all parameters of electrochemical potential equation [41] – voltage, concentration, and temperature, since temperature now becomes an ultra-local variable.

The questions related to the ultra-local thermodynamic effects hiding behind the curtains of the highly probable Joule heat release on open ionic channels are of primary interest, as they are currently dramatically underestimated [31]. The heat conductivity coefficient of intracellular media is 5 to 7 times smaller than that of water [42], [43], therefore one can imagine the presence of steep nano-/microscale temperature gradients both for artificial micro-/nanoscale heat sources, as well as for probable endogenous intracellular “hot spots” such as mitochondria, ion pumps, and ion channels. For example, an open ion channel operating under electrical potential on cellular membranes is one of the most intriguing cases of suspected intracellular ultra-local heat sources. Remarkably, up to 10’s °C temperature rise due to Joule heat release was calculated using nanoscale hydrodynamic modeling of ion velocities distributions accelerating in the ion channels [31], where the typical transmembrane electric field strength on the order of −70 mV/7 nm corresponds to ~10 MV/m, which is comparable to the typical electric field strength in particle accelerators [44]. These data are in line with the hypothesis of the thermal signaling concept [45], which involves endogenous ultra-local thermal control of spatially localized intracellular physical and biochemical processes by nanoscale intracellular spot heat sources, referred to as “nanoscale boilers” [31].

Therefore, the existing paradigm neglecting the ultra-local effects of intrinsic cellular thermodynamics must be updated. Currently, temperature is typically viewed as a macroscale parameter that depends solely on the macroscale heat production and heat conduction in the tissue or body of a multicellular organism, or as an influence of external, usually slowly changing, environmental temperature. The updated view should also consider temperature as an intracellular ultra-local variable once we get to the micro/nanoscale. However, on the way to this new paradigm, at least two serious problems existed. The first issue is the clear deficiency of simple, reliable experimental methods for micro- and nanoscale temperature measurements and control in aqueous media near and inside living cells, as discussed in [1], [4], [18], [21]. The second concern is the widespread doubt regarding the validity of using correct temperatures in micro- and nanoscale aqueous volumes. To overcome this notion, one needs to remember that the fundamental temperature definition is as an equilibrium macroscopic intensive thermodynamic parameter [16] assigned to the macroscopic thermodynamic systems (namely systems with macroscopic amounts of molecules (particles), but not necessarily macroscopic in size as discussed in [5].

Albert Einstein discussed the ultimate importance of different aspects of validity limits in classical thermodynamics in a lecture he delivered at the University of Bern in 1908 [46]. It is a non-trivial question - where indeed can we theoretically expect approaching spatial and temporal limits for the ultra-local temperature definition inside a living cell? Namely, when will we still have the right to define different temperatures of neighboring intracellular nano/microscale aqueous volumes, given the fact that there are multiple potential ultra-local nanoscale heat sources in living cells? How do we justify acceptable theoretical correctness for the definition of such stationary or transient ultra-local temperature gradients? As it was discussed in [21], the definition of the size of an elementary δV of aqueous media (which should itself contain a macroscopic number of molecules) reflects the physical limit of the time resolution of the description of the temperature gradients, which cannot be better than the time of local relaxation. For example, if we choose a space resolution of δV = 1 cubic micron, the time resolution cannot be better than 10 microseconds.

The same logic applies to justifying the correctness of the temperature profile clamp method at the nano-/microscale by the DHT device described in this paper. The ultra-local nano-/microscale thermodynamic variable – temperature – is a macroscopic equilibrium intensive parameter, as it first appeared correctly in Maxwell’s statistical distribution of molecules by speeds. Therefore, temperature can be freely used (controlled and measured) in the aqueous media including the living cell down to characteristic dimensions of 10 nanometers (with a corresponding limit of time resolution) as it is currently accepted for the other parameters of the electrochemical potential equation [41] – electrical potential and ionic concentrations. Simple physical estimations in line with [21] justify that there are no theoretical problems in defining the local temperature for an elementary intracellular volume down to ~100 nm^3^. The problems begin for such ultra-local aqueous systems when approaching 10 nm^3^, the characteristic length scale of the cell membrane thickness. Such yoctolitre volume (10^−24^ L) contains around *N* = 33×10^3^ molecules of water giving a rise to spontaneous relative fluctuations in the thermodynamic parameters on the order of N^−1/2^ (~0.006), resulting in 1.8 K temperature fluctuations [38] that will further grow to ~600K once the volume of aqueous medium is reduced to 1 nm^3^.

## Conclusions

In conclusion, we presented a new methodology for a micro-/nanoscale temperature controller in aqueous media, enabled by the Diamond Heater-Thermometer in a fully optical fiber-based configuration, combined with the patch clamp technique. Using this approach, we have demonstrated applications for the local, fast, and reproducible thermal modulation of ionic current from voltage-gated Na_v_1.5 sodium channels present in cardiomyocytes and overexpressed in heterologous expression systems. The presented approach of manipulating ultra-local intracellular temperatures down to the nanoscale can uncover previously inaccessible effects in various physiological intracellular processes related to the endogenous nanoscale heat sources (e.g., heat release in the open ion channels, ionic pumps, uncoupled mitochondria, and local calcium release processes). Moreover, thanks to the configuration of the demonstrated fiber-coupled Diamond Heater-Thermometer, its ease of use and versatility, such a technique can become part of a standard toolbox for electrophysiology, calcium imaging, and experimental life sciences in general.

## Materials & Methods

### Diamond Heater-Thermometer (DHT) device

A fully fiber-coupled version of the DHT device was used. It consisted of a glass pipette with a micron-scale diamond with embedded temperature-sensing silicon-vacancy (SiV) centers placed at its tip. The diamond particle was connected to a tapered optical fiber serving both to guide the excitation light and to collect the fluorescence emitted by the SiV centers. The excitation was provided by a fiber-coupled laser (520 nm, CNI lasers), and a fiber-coupled spectrometer (ATP5200, OptoSky Photonics) was used to detect the fluorescence peak. The fibers were connected using a fiber splitter (Thorlabs). Such a DHT configuration can be easily added to any optical microscope, like the Zeiss Axio Z1 microscope used in this work.

### Cell line preparation

Human embryonic kidney (HEK293) cells stably expressing the human voltage-gated sodium channel Na_v_1.5 were cultured in DMEM (Gibco, Basel, Switzerland) supplemented with 10% FBS, 0.5% penicillin, Zeocin (200 µg/mL), and streptomycin (10,000 U/mL) at 37 °C in a 5% CO_2_ incubator.

### Isolation of mouse ventricular myocytes

All animal experiments were performed according to the Swiss Federal Animal Protection Law and approved by the Cantonal Veterinary Administration, Bern. This investigation conforms to the Guide for the Care and Use of Laboratory Animals, published by the US National Institutes of Health (NIH publication no. 85-23, revised 1996).

Single cardiomyocytes were isolated according to a modified procedure of established enzymatic methods. Briefly, mice were deeply anesthetized using a ketamine/xylazine mix (200/20 mg/kg body weight) via intraperitoneal injection. After losing reflexes, hearts were rapidly excised, cannulated, and mounted on a Langendorff column for retrograde perfusion at 37 °C. Hearts were rinsed free of blood with a nominally Ca^2+^-free solution containing (in mM): 135 NaCl, 4 KCl, 1.2 MgCl_2_, 1.2 NaH_2_PO_4_, 10 HEPES, 11 glucose, pH 7.4 (NaOH adjusted), and subsequently perfused by a solution supplemented with 50 µM Ca^2+^ and collagenase type II (0.5 mg/mL, Worthington, Allschwil, Switzerland) till achieved digestion. Following digestion, the atria were removed, and the ventricles were transferred to a nominally Ca2+-free solution, supplemented with 100 µM Ca2+, and cut into small pieces. Single cardiac myocytes were liberated by gentle trituration of the digested ventricular tissue (using a 1 mL pipette with a wide-bore tip) and filtered through a 100-180 µm nylon mesh. Ventricular mouse cardiomyocytes were used after an extracellular calcium increase procedure to avoid calcium overload when extracellular solutions were applied in electrophysiology experiments.

### Whole-cell electrophysiology

Sodium currents (I_Na_) were recorded in the whole-cell configuration using a VE-2 amplifier (Alembic Instrument, USA). Borosilicate glass pipettes were pulled to a series resistance of ~2 MΩ. pClamp software, version 8 (Axon Instruments, Union City, CA, USA) was used for recordings. Data were analyzed using pClamp software, version 8 (Axon Instruments), and OriginPro, version 7.5 (OriginLab Corp., Northampton, MA, USA).

I_Na_ in HEK293 cells was carried out using an internal solution containing (in mM) CsCl 60, Cs-aspartate 70, CaCl_2_ 1, MgCl_2_ 1, HEPES 10, EGTA 11, and Na_2_ATP 5 (pH was adjusted to 7.2 with CsOH). The cells were bathed in a solution containing (in mM) NaCl 50, N-Methyl-D-glucamine (NMDG)-Cl 80, CaCl_2_ 2, MgCl_2_ 1.2, CsCl 5, HEPES 10, and glucose 5 (pH was adjusted to 7.4 with CsOH).

I_Na_ in cardiomyocytes was carried out using an internal solution containing (in mM) CsCl 60, Cs-aspartate 70, CaCl_2_ 1, MgCl_2_ 1, HEPES 10, EGTA 11, and Na_2_ATP 5 (pH was adjusted to 7.2 with CsOH). Cardiomyocytes were bathed in a solution containing (in mM) NaCl 5, NMDG-Cl 125, CaCl_2_ 2, MgCl_2_ 1.2, CsCl 5, HEPES 10, and glucose 5 (pH was adjusted to 7.4 with CsOH). Nifedipine (10 µM) and cobalt chloride (CoCl_2_) (10 µM) were added to the extracellular solution to inhibit calcium currents.

For cardiac action potential (AP) recordings, cardiomyocytes were bathed in a solution containing (in mM) NaCl 140, KCl 5.4, CaCl_2_ 1.8, MgCl_2_ 1.2, HEPES 10, and glucose 5 (pH was adjusted to 7.4 with NaOH). Cardiomyocytes were initially voltage clamped (holding potential −80 mV) and dialyzed with an internal solution containing (in mM) KCl 120, CaCl_2_ 1.5, MgCl_2_ 5.5, Na_2_ATP 5, K_2_-EGTA 5, and HEPES 10 (pH was adjusted to 7.2 with KOH). APs were elicited at 0.5 Hz with rectangular pulses (5 ms at 125% threshold) in current-clamp mode. Elicited APs were allowed to stabilize before one or more sequences of ~1 minute each were acquired from each cell. AP recordings were digitized at a sampling frequency of 20 kHz. Electrophysiological data were analyzed offline, where the resting membrane potential, time to maximum amplitude and AP durations (APD) at 30, 50, and 90% repolarization were averaged from each sequence of APs.

### Statistical analysis

Data are represented as means ± S.E.M. Statistical analyses were performed using Prism7 GraphPad^™^ software. A Wilcoxon signed-rank test was used to compare two groups due to the small sample size. *p* < 0.05 was considered significant.

## Acknowledgments

We thank Dr. Sarah Vermij for her useful comments and proofreading on this manuscript.

