## Supplementary figures for "ll-optical Diamond Heater-Thermometer enables versatile and reliable thermal modulation of ion channels at the single-cell level"

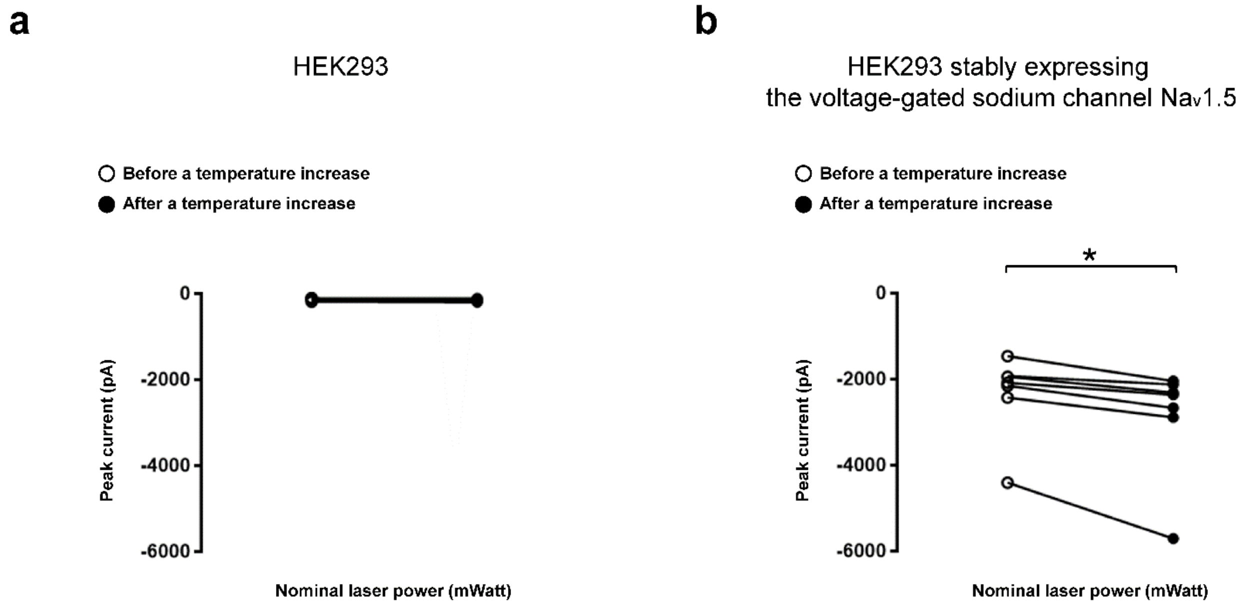


Figure S1. Only the voltage-gated sodium channels Nav1.5 are modulated by the local heat pulse: a) wild-type cells; b) cells stably expressing the voltage-gated sodium channels Nav1.5.

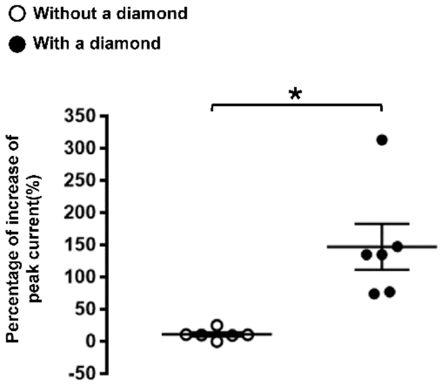


Figure S2. Laser illumination does not affect the biophysical properties of the Nav1.5 channel, as shown by the difference in the peak current increase when a pipette without (white circle) and with (black circle) a diamond is used at a laser power of 40 mW (*; $p < 0.05$).
